# Genomic epidemiology of the 2017-2023 outbreak of *Mycoplasma bovis* sequence type ST21 in New Zealand

**DOI:** 10.64898/2026.04.07.717125

**Authors:** Nigel P French, Amy Burroughs, Barbara Binney, Samuel J Bloomfield, Simon M Firestone, Jonathan Foxwell, Edna Gias, Kate Sawford, Mary van Andel, David Welch, Patrick Biggs

## Abstract

*Mycoplasma bovis* was first detected in cattle in New Zealand in 2017, prompting an eradication programme that incorporated extensive surveillance and a test-and-cull policy. Genome sequence data and phylodynamic models were used to inform decision making throughout the eradication programme. Isolates from 697 cattle on 126 farms were collected and sequenced between July 2017 and December 2023. Phylodynamic models were used to estimate the time of most recent common ancestor, the effective reproduction number (R_eff_) and effective population size, and long-range and local between-farm transmission dynamics. The analysis revealed the dramatic impact of movement restrictions and culling up to early 2020, with a sharp reduction in the R_eff_ to less than 1 in 2018/9 and the extinction of two of three major lineages in 2020. This was followed by three-years of residual infection in farms in the South Island, associated with persistent infection of a large feedlot farm and nearby farms. The comprehensive dataset of genomic and epidemiological data provided a unique opportunity to study the dynamics of a country-wide outbreak of a single-host pathogen from first detection to potential eradication, underlining the utility of integrated genomic surveillance during an outbreak response.

**Author summary:** The economically important cattle pathogen, *Mycoplasma bovis*, was first detected in New Zealand in 2017. This led to a large-scale, successful control programme aimed at eradication of the pathogen. The decision to undertake an eradication programme was informed by initial analyses of whole genome sequences from isolates collected as part of the surveillance programme. The analysis showed that the bacteria had entered New Zealand relatively recently and was unlikely to be widespread. Over the subsequent years, genome sequencing and modelling of transmission dynamics informed important policy decisions made by the New Zealand Government and the cattle industry, and helped to monitor progress of the eradication programme. The impact of the detection, movement control and culling programme was profound, with sharp reductions in transmission between 2018 and 2020. This was followed by a long tail of localised infection in the South Island, involving transmission from a large feedlot farm. Provisional eradication was achieved after depopulation of this feedlot. This analysis highlights the role of genomic surveillance and modelling to inform decision making during an infectious disease outbreak.

## Introduction

*Mycoplasma bovis* is a globally important cattle pathogen, with major economic and welfare impacts in cattle industries around the world [1]. After its initial discovery as a cause of mastitis in the USA in 1961 [2], it has been reported in Australia, Europe, South America, Japan, Ireland, China, India and Africa [1, 3–6]. New Zealand was considered free of *M. bovis* until it was first reported in the South Island in 2017 [7, 8].

*M. bovis* is associated with several clinical diseases, including pneumonia, mastitis, arthritis, genital disorders and keratoconjunctivitis and otitis media [1, 9, 10]. Animal-to-animal transmission occurs via colostrum, milk, semen, airborne droplets, intrauterine transmission [4] and potentially via contaminated environments [11] with farm-to-farm transmission occurring via movements of cattle, colostrum, milk and semen. Treatment, control and prevention are increasingly difficult in the face of antimicrobial resistance and the lack of a commercially available and effective vaccine [12–14]. New Zealand’s economy is highly dependent on agriculture, and the dairy and beef industries are major contributors to the country’s total export trade in commodity goods – accounting for 29% and 6% of the value of total goods exported in 2024 respectively [15]. The detection of *M. bovis* in New Zealand on 22 July 2017, and the potential for rapid spread in a large and previously unexposed cattle population, was therefore viewed as a major risk to the country’s cattle industry and the economy as a whole.

In response to this threat an industry-wide ‘*Mycoplasma bovis* Eradication Program’ was implemented in May 2018 [8, 16]. The program involved multiple surveillance streams, including tracing of cattle and milk movements between farms, extensive testing of bulk milk samples [17], and culling of both dairy and beef herds confirmed as infected with *M. bovis* [8, 16]. Following depopulation of infected herds, the premises were cleaned and disinfected and a stand down period of 60 days was mandated for dairy farms prior to repopulation. Whole genome sequencing was deployed early in the outbreak to characterise the strains detected and assess their diversity [18]. The simultaneous collection of genome sequence and epidemiological data provided a unique opportunity to use phylodynamic methods [19] to assess the time of origin of the New Zealand isolates, and the potential number of incursions. As the outbreak progressed, genomic data were also used to support investigations into likely transmission pathways between infected farms.

A range of tools were used throughout the outbreak to interpret genomic and epidemiological data and to contribute to decision-making by the response team. This included the use of visualisation tools such as Nextstrain [20] and Microreact [21] to overlay the phylogeny onto spatial and other data, and Bayesian phylodynamic models [22–24]. Phylodynamic models were used to infer the time of incursion of *M. bovis* into New Zealand, how the effective reproduction number changed over time, and both long range movements and local networks of transmission between infected farms. In this study we describe the genomic epidemiology of the entire outbreak, from the first isolates recovered in 2017 to the most recent in December 2023 and include examples of outputs generated during the outbreak as sequence data became available.

## Methods

A schematic of the methods used to analyse genomic data and inform decision making throughout the outbreak is provided in Figure 1.

**Figure 1.**
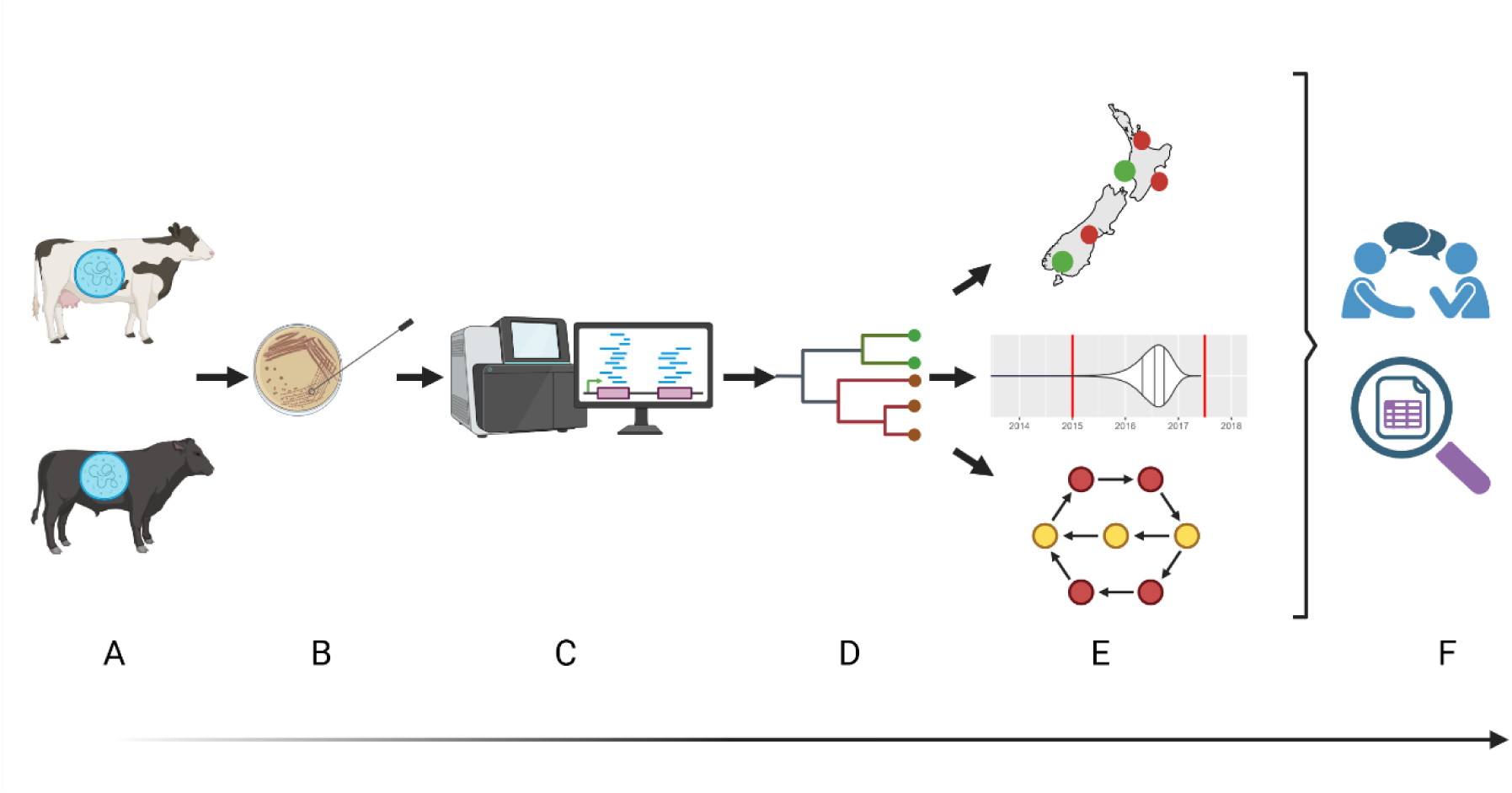
Steps in the application of genomic surveillance for the investigation of the *Mycoplasma bovis* outbreak in Aotearoa New Zealand, from detection of infected farms to informing decision making. A: Farm identified as infected through surveillance. B: Samples from infected cattle collected and cultured to isolate colonies of *M. bovis*. C: DNA extracted and analysed using Illumina and PacBio sequencing. D: Read data analysed using bioinformatics tools to generate data for phylogenetic and phylodynamic analysis. E: Data visualised using Nextstrain and Microreact, and phylodynamic modelling used to determine the date of common ancestor and between farm transmission probabilities. F: Outputs are discussed with decision makers in light of other data, and further targeted analyses conducted.

### Sampling

Details of sample collection and *M. bovis* culture are provided by Binney, Gias (18). Further information about the outbreak response and detection of infected farms is also provided by Jordan, Sadler (8), [16]. In brief, samples from real-time PCR positive cattle detected during the surveillance programme were cultured for *M. bovis*. Bacterial isolation was successful from 697 cattle, across 126 farms, including nasal and oropharyngeal cavities, pharyngeal tonsils, synovial fluid, lung tissue and milk. The 126 farms represented all farms from which *M. bovis* could be isolated and sequenced, out of a total of 282 farms confirmed as infected (Table S1). Samples were inoculated in Friis broth (FB) and inoculated onto Friis agar (FA) [25] incubated at 37°C, 5% CO_2_. Single colonies were then isolated and inoculated in FB, and DNA extracted after 3–4 days.

### Whole genome sequencing

The processes used for DNA extraction, *M. bovis* confirmation, quality control and short read Illumina sequencing are described by Binney et al. (2025). In brief, DNA was extracted using the QIAamp DNA Mini Kit (QIAGEN, Hillden, Germany), DNA concentration measured by Qubit dsDNA High Sensitivity (HS) Assay kit (Thermo Fisher Scientific, Eugene, Oregon, United States) and quality assessed by NanoDrop spectrophotometer (Thermo Fisher Scientific, Wilmington, Delaware, United States). The VetMax^TM^ M. bovis kit (Life Technologies, Carlsbad, CA, United States) was used to confirm the presence of *M. bovis* DNA. DNA libraries were prepared using the Nextera XT DNA Library Preparation Kit (Illumina, San Diego, CA, United States) following the manufacturer’s protocol, and the concentrations and quality of the libraries were assessed using the Qubit dsDNA HS Assay kit and HS DNA kit on an Agilent 2100 Bioanalyzer or the HS D5000 ScreenTape kit on the 4200 TapeStation (Agilent Technologies, Waldbronn, Germany). Sequencing was performed on an Illumina MiSeq using the MiSeq reagent Kit v2 (500 cycles) (Illumina).

A reference isolate (NCBI CP192245.1) was generated using long-read sequencing (Pacific Biosciences, Inc., RS II platform) using P6-C4 chemistry according to the 20 kb Template Preparation and the BluePippin DNA Size Selection system protocol (Pacific Biosciences, Inc.) as described in Binney, Gias (18).

### Bioinformatic analyses

#### Quality checks and genome assembly

The pipeline used for sequence quality checks and genome assembly is described in detail in Binney, Gias (18). In brief, quality checks were carried out using phyloFLASH v3.31b1 [26] after trimming sequences using Trimmomatic v0.39 [27]. The output files were evaluated by FastQC v0.11.9 [28] and reviewed using MultiQC v1.8 [29]. The Nullarbor v2.0.20191013 pipeline assembled all the sequences into draft genomes [30]. Draft genome assemblies produced by SKESA v2.4.0 were used [31].

#### Genome datasets

Isolates from 697 cattle on 126 farms were collected and sequenced between July 2017 and December 2023. Single isolates were randomly selected from each animal and each farm, and these samples are referred to as the animal-level and farm-level datasets respectively. Subsets of these datasets were used for analyses covering different time periods and different lineages.

#### Single nucleotide polymorphism identification

Single nucleotide polymorphisms (SNPs) were identified using the Snippy v2.6 pipeline (https://github.com/tseeman/snippy). Snippy used the Burrows-Wheelers Aligner [32] and SAMtools [33] to align reads from different isolates to the New Zealand ST21 reference genome (NCBI CP192245.1) and FreeBayes [34] to identify variants among the alignments. Gubbins [35] was used to remove regions of the genome identified as recombinant.

### Phylogenetic analysis

Support for a temporal signature (i.e. evidence for a strict molecular clock) in the animal- and farm-level datasets was assessed by estimating the correlation between the optimal phylogenetic tree branch length and the sampling dates. Models comparing the actual data against a simulated tree, imposing a single sample date, were used to further test for evidence of a temporal signature [36].

The SNPs were imported into BEAUti 2.5 to create an Extensive Markup Language (.xml) file for BEAST 2.7.8 [22]. The *M. bovis* reference genome (NCBI CP192245.1; 1,064,193 bp) was used to estimate the number of invariant sites which were added as constant site weights to the .xml file.

### Population structure – clade analysis and time tree

Population clustering of isolates sampled from individual cattle was explored using the rhierBAPS algorithm in R [37, 38]. This is a Bayesian method for hierarchically clustering sequence data to reveal nested structures. The maximum number of populations was assumed to be 20, and the maximum depth of hierarchical search was set at 2. Only the level 1 clades are reported. This analysis was repeated throughout the outbreak as new sequence data became available. In this study we report the clade structure from the full animal-level dataset.

A maximum clade credibility (MCC) tree was generated from the posterior distribution of trees output from a BEAST model of the full animal-level dataset. For this model we used an HKY substitution model, a strict clock and a Bayesian Integrated Coalescent Epoch PlotS (BICEPS) tree prior [39].

### Time of most recent common ancestor (tMRCA), effective reproduction number (R_eff_) and effective population (N_e_) size estimation

The tMRCA was repeatedly estimated from the animal- and farm-level datasets throughout the outbreak as new data became available. Multiple substitution, molecular clock and tree models were trialled for each estimate of the tMRCA. Nested sampling (NS) [40] was used to select models.

Combinations of molecular clock including strict and uncorrelated relaxed clocks [41] and tree priors including the birth-death skyline (BD-SKY) [24] and Extended Bayesian Skyline (EBS) [42] were trialled. The .xml files were run in BEAST for 50 million steps, three times with different starting seeds, before LogCombiner was used to combine the runs with a 10% burn-in.

The tMRCA was estimated from the final farm-level dataset (N=126) using the site model averaging bModelTest clock prior [43], uncorrelated relaxed clock [41] and BD-SKY [24]. The tMRCA was estimated from the final animal-level dataset (N=697) using the tree height estimate from the BICEPS model described above [39].

A uniform prior was placed on strict clock substitution rates with an upper bound of 10^-4^ substitutions site^-1^ year^-1^ and a lower bound of 10^-8^ substitutions site^-1^ year^-1^ [19]. LogCombiner was used to combine the runs with a 10% burn-in. Tracer v1.6 [44] was used to visualise the results. TreeAnnotator 2.6 was used to generate a MCC tree.

The Tree Model Adequacy (TMA) package in BEAST2 [45] was used to check the fit of the MCC tree produced under constant and exponential coalescent, and BD-SKY models. The model was run for 10 million steps and 100 trees. The following statistics were analysed: tree height, time of maximum lineages, slope ratio between the tree origin and the maximum number of lineages, the Colless tree imbalance index and the external/internal branch ratio.

The effective reproduction number (R_eff_) was calculated from the BD-SKY model [24] applied to the final farm-level dataset available in December 2023. R_eff_ was calculated for ten time-intervals. The effective population size (N_e_) was estimated directly from the BICEPS model described above.

### Structured coalescent modelling

#### Farm to farm transmission

Transmission between farms was investigated using the Structured COalescent Transmission Tree Inference model (SCOTTI) as described by De Maio, Wu (23). Models were compiled using the ‘SCOTTI_generate_xml.py’ script [46] and run in BEAST2 [22]. Animal-level datasets were used in this analysis, using isolates from all infected farms with available sequences. Subsequent analyses focused solely on lineage ‘a’ (RB clades 2, 4 and 5). These analyses were continually updated as the outbreak progressed. Two of the three initial lineages (‘b’ and ‘c’) were not detected after 2020 and assumed to be extinct. Two analyses are presented in this study; the dataset of all lineages up to December 2020 and the dataset of lineage ‘a’ isolates throughout the outbreak to December 2023.

Each infected farm was considered a distinct population (deme) and was assigned a period of infectiousness, which started with the estimated date on which the farm became infected. This was based on epidemiological estimations of when infection was first introduced to a farm considering traced movements from known infected farms, farming practices, onset of clinical signs (if present) and laboratory data. The end of the infectious period was the later of the following two dates: the date when movement controls were imposed on the farm and the date of sampling of the last isolate from each infected farm submitted for genome sequencing. The model was run in triplicate for 50M iterations and the output was processed to determine the transmission network and the probability of direct, indirect and total transmission between each pair of infected farms, using a modified version of the python script ‘Make_transmission_tree_alternative.py’ [46]. Convergence was assessed in Tracer (v1.644), [47], and all effective sample sizes were >300.

To assess the sensitivity of the model to the onset date and period of infectiousness the analysis was repeated three times, using the onset of infectiousness dates estimated from epidemiological estimates made using data available in 2023 and 2025, and a date of one month prior to the imposition of movement restrictions. The latter was considered the most extreme plausible estimate of date of onset of infectiousness and imposed narrower estimates of the infectious period (range 0 – 5.1 years shorter).

#### *National-level dispersal of* M. bovis *following incursion*

The long-range dispersal of *M. bovis* from incursion to the end of 2020 was inferred using the Marginal Approximation of the Structured Coalescent (MASCOT) model (v3.0.7) [48] including time-varying predictors of *N_e_* using MASCOT-GLM [49]. We were primarily interested in migration (transmission events) between the North and South Island using a two-deme model. *N_e_* predictors were the weekly prevalence of infected farms in each island. All other parameters and settings were identical to the BICEPS model described above. The posterior distribution of trees was compared using MCC and conditional clade distribution (CCD-0) trees [50].

To further investigate evidence for transmission to and from a large feedlot farm located in the South Island, a three deme model was constructed comprising the feedlot, other South Island farms and North Island farms.

## Results

### Datasets

A total of 1026 isolates from 697 cattle on 126 farms were genome sequenced as part of the investigation up to December 2023. This represented isolates from 45% of all infected and depopulated farms (126/282). The number of cattle from which isolates were sequenced per farm ranged from 1-99, with the largest number of isolates selected from a large feedlot farm that was persistently infected over a four-year period, and one of the last farms to be depopulated (in December 2022). A random sample of single isolates from each animal (animal-level dataset N=697) and each farm (farm-level dataset N=126) were selected for the analyses described below.

### Population structure and epidemic timeline

The animal-level dataset was used to describe the population structure and dynamics over the course of the outbreak. The distribution of SNP distances for this dataset is shown in Supplementary Figure S1; the root-to-tip analysis correlation coefficient was 0.82, the R^2^ value was 0.67 and there was stronger support for a model with tip dates corresponding to the date of isolation than a model with uniform simulated dates.

Figure 2A shows the time course of the outbreak, with the BEAST time tree (using the BICEPS model) of animal-level isolates (N=697) revealing the population structure with tip dates corresponding to the date of *M. bovis* isolation. The time series of all movement restrictions, depopulations and active case farms are plotted on the same scale (Figures 2B, 2C and 2D respectively). Most confirmed infected farms were identified and depopulated between 2018 and mid-2020.

**Figure 2.**
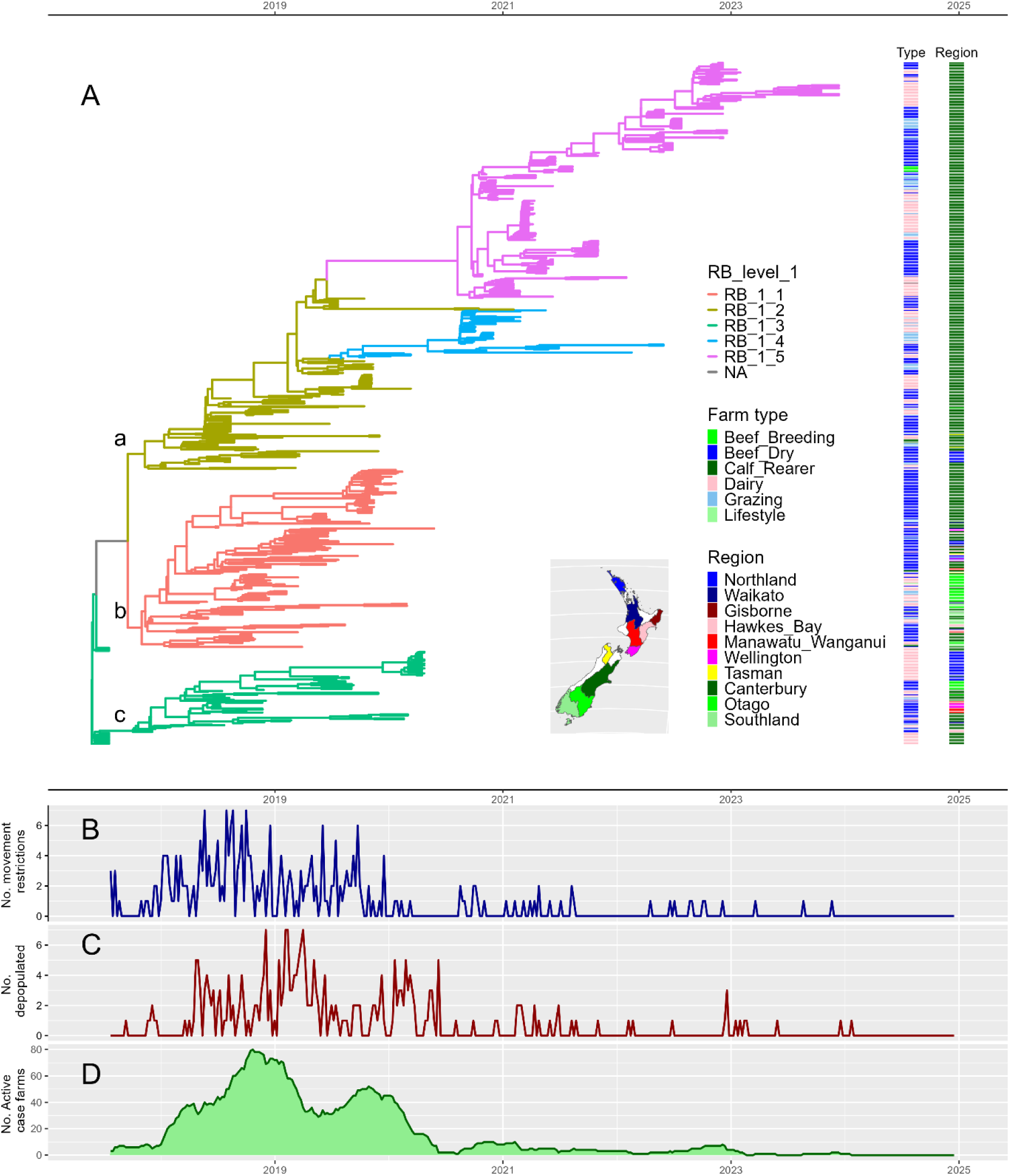
Time tree of 697 randomly selected isolates from infected cattle and the time series of key measures of herd-level infection and intervention plotted on the same x-axis. Figure 2A shows the MCC tree from the BICEPS model where the tips of the tree correspond to the time of isolation of *M. bovis* on the x-axis. Tree lineages are coloured by RB level and the three main lineages evident in 2017 are marked a, b and c. Posterior support for these lineages was >0.9. The heatmaps to the right of the tree show the farm type (right), and the region of New Zealand the isolates were sampled from (far right, with key in inset map). The three lower plots show the time series of weekly incidence of the number of farms with movement restrictions imposed (Figure 2B), the incidence of the number of farms depopulated (Figure 2C) and the prevalence of detected, actively infected farms (Figure 2D).

Initial analyses conducted between 2018 and 2020 provided evidence of the emergence of three lineages in ∼2017, designated ‘a’, ’b’ and ‘c’ with posterior support of 0.99, 0.99 and 0.98 respectively. As the outbreak progressed lineage ‘a’ diverged into three clades, designated RB_1_2, RB_1_4 and RB_1_5 (RB stands for rheirBAPS). From 2021 onwards, only clades RB_1_4 and RB_1_5 were identified, and the outbreak was confined to the Canterbury Region in the South Island. Multiple farm types were infected throughout the outbreak (Figure 2A).

### Time of most recent common ancestor (tMRCA)

The tMRCA was estimated throughout the outbreak to provide information on the most likely time of incursion for the programme. The choice of substitution, clock and tree models varied over time.

However, the estimated tMRCA remained relatively constant from the first estimation in 2019 to the final estimate in 2023, with most posterior distributions estimated to be between 2015 and mid-2017 (Supplementary Figure 2).

The final median estimated tMRCA was April 2016 (95% highest posterior density (HPD) interval: August 2015 - October 2016) for the farm-level dataset and May 2017 (95% HPD interval: April 2017-June 2017) for the animal-level dataset.

### Effective reproduction number (R_eff_) and effective population size (N_e_)

The change in herd-level *R_eff_* over time was estimated from the full farm-level dataset and the BD-SKY model used for the final estimate of the tMRCA. This analysis showed a marked decline in the *R_eff_* from ∼2.5 to 0.5 from mid-2018 onwards, after the onset of movement restrictions on many farms and culling of cattle on infected farms (Figure 3A). This is plotted with the *N_e,_* estimated from the animal-level dataset using the BICEPS model (Figure 3B). *N_e_* is influenced by the total pathogen population and pathogen diversity, both of which are affected by interventions that reduce the number of infected individuals and induce evolutionary bottlenecks. *N_e_* increased from the tMRCA to 2019 and then decreased to mid-2020, before a smaller increase in 2021. This trend is broadly consistent with the number of active case farms shown in Figure 2.

**Figure 3.**
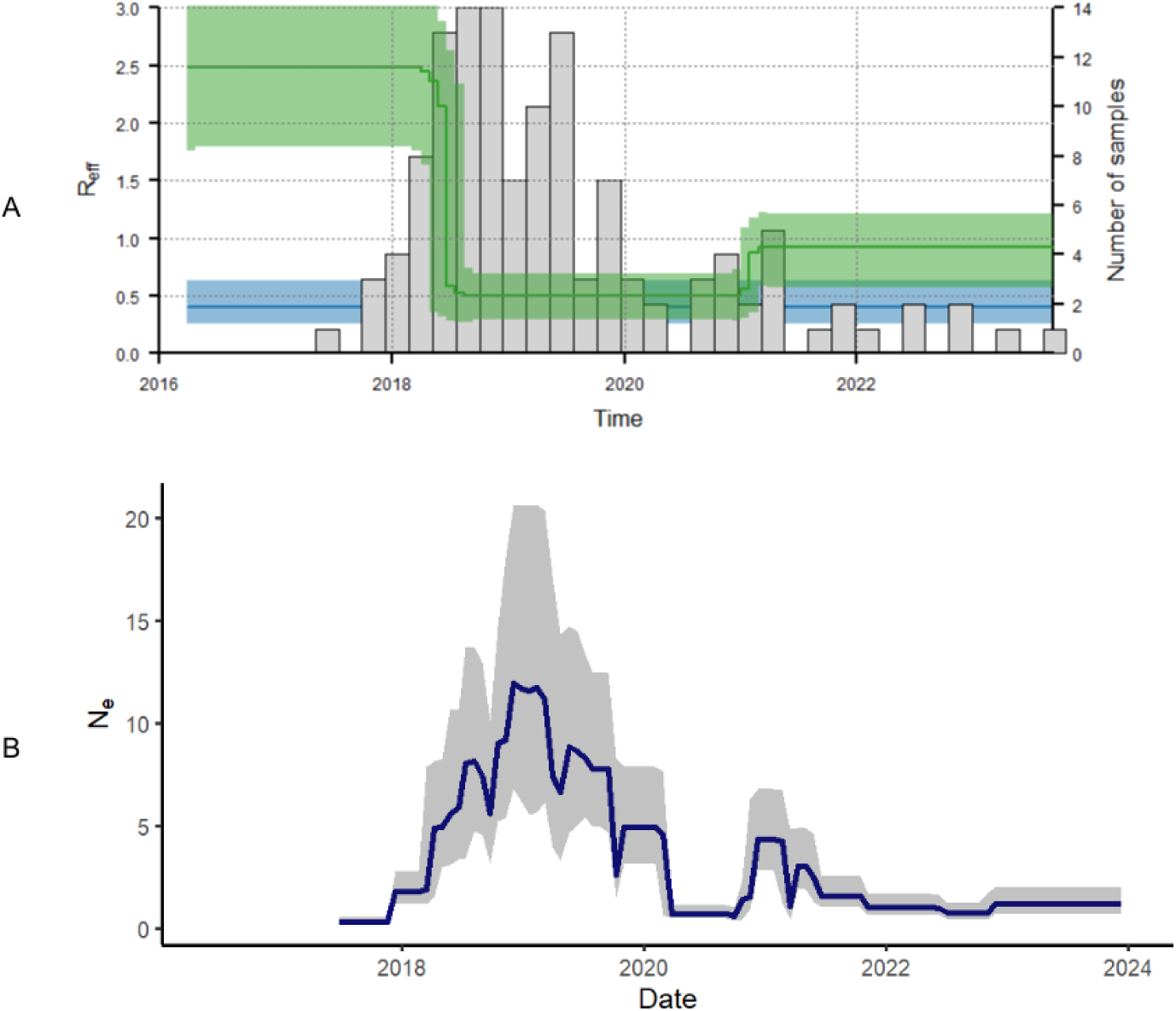
A: Analysis of a BD-SKY model of randomly selected isolates from 126 farms infected with *M. bovis* (single isolate per farm) showing the estimated effective reproduction number, *R_eff_*, (left y-axis, green line with shaded 95% HPD), number of samples (grey bars) and estimated rate at which infected farms become uninfectious through isolation (e.g. movement restrictions) or culling (blue line with shaded 95% Credible Intervals and the same scale as right y-axis showing the number of samples). B: Effective population size, *N_e_*, (blue line with shaded 95% HPD) estimated from animal-level samples and the MCC tree estimated from the BICEPS model illustrated in Figure 2.

### SCOTTI analysis

Figure 4A shows the estimated between-farm probabilities of direct transmission using data up to December 2020. Only probabilities greater than 0.4 are shown. There is evidence of localised transmission between dairy and non-dairy farms and multiple long-range transmission events, mostly between the South and North Island. Four farms are associated with multiple transmission events (six or more), including three dairy and one beef farm located in the South Island.

Figure 4B shows the estimated network of direct transmission of persistent lineage ‘a’ throughout the outbreak (i.e. RB clades 2, 4 and 5 in Figure 2A). This lineage was primarily located in the Canterbury region in the South Island. One large beef feedlot in Canterbury, that was persistently infected for at least four years between August 2018 and December 2022, was associated with multiple transmission events to and from other local farms, all in the Canterbury region. The direction of transmission indicates both inward and onward transmission. Supplementary Figure S3 is the equivalent network of direct transmission using the alternative date of onset of infectiousness of one month prior to the imposition of movement restrictions. The pattern of transmission is similar and provides further support for the role of the feedlot farm in onward transmission, with multiple farms predicted to be infected with genomic descendants of lineages circulating in the feedlot.

The key showing region colours is included (Figure 4C). Single randomly selected isolates from all available individual animals were included in the analyses and only direct transmission probabilities >0.4 are shown. D=dairy, G=grazing, B=beef, Br=beef breeding, L=lifestyle.

**Figure 4.**
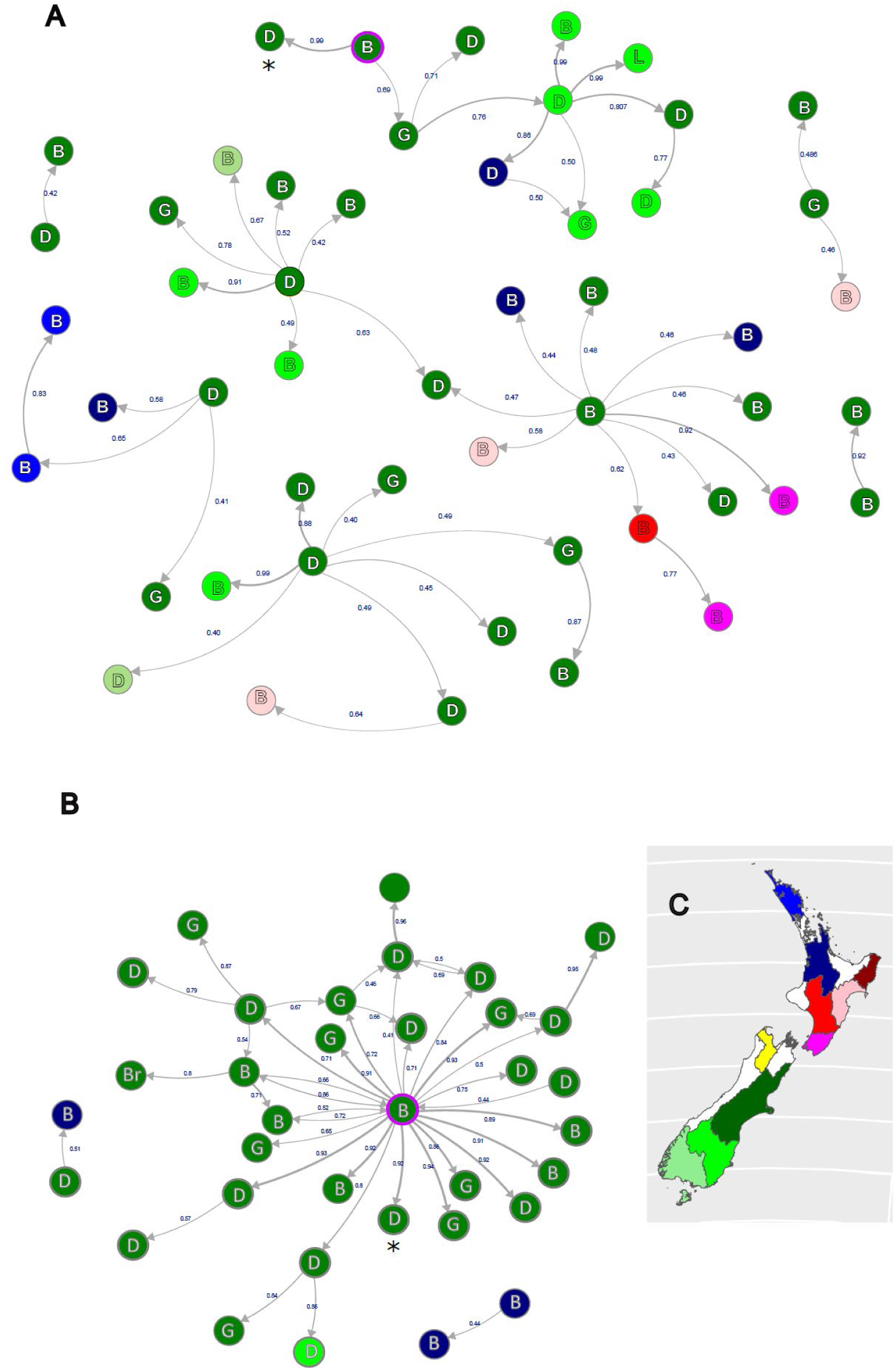
Examples of network diagrams showing between-farm direct transmission probabilities estimated using the SCOTTI model, from all farms with genome sequenced isolates up to December 2020 (Figure 4A) and all lineage ‘a’ isolates (RB levels 2, 4 and 5) up to December 2023 (Figure 4B). Some nodes are common to both networks. For example, the central node in Figure 4B labelled B with a purple border is the feedlot from Canterbury referred to in supplementary Figure S4, and this node is also evident with a purple border in the top of Figure 4A. In both plots the feedlot is connected to a dairy farm with probability >0.9, marked with an asterisk.

### *Long range* M. bovis *transmission events*

Analysis of ‘migration’ between the North and South Island using a 2-deme MASCOT GLM model provided evidence of multiple transmission events in both directions (median estimated 12 migrations from the South to North Island and 5 migrations from the North to South Island (Figure 5).

**Figure 5.**
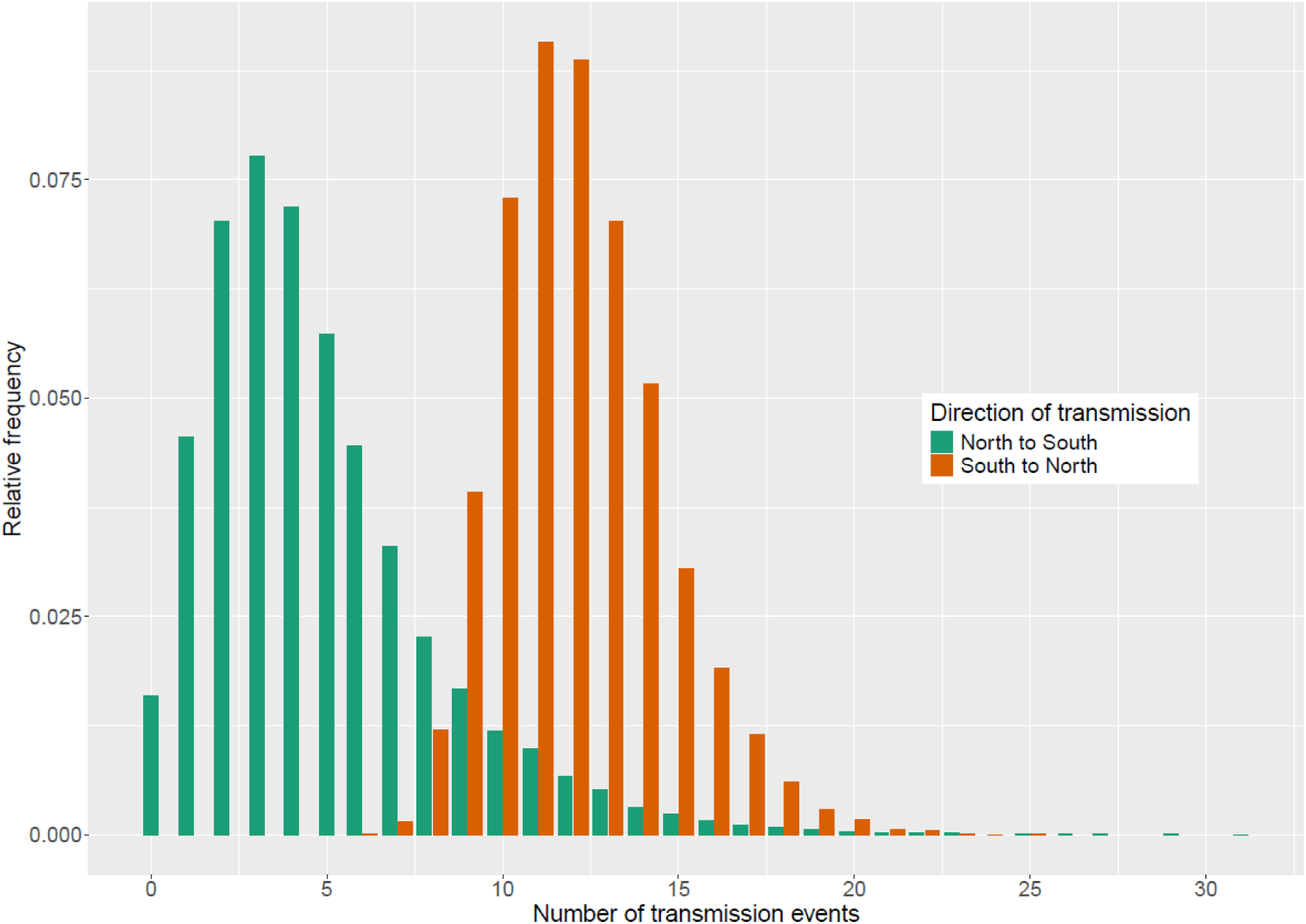
Posterior distributions of estimated between-island *M. bovis* transmission events from a 2 deme marginal structured coalescent approximation model, MASCOT GLM [48, 49], in BEAST2 [22].

The MCC tree shows the most likely ancestral location for internal nodes within the tree (Figure 6). There was no evidence of migrations between the two islands after the end of 2019 despite new incident cases being detected in both the North and South Island in 2020 (Figures 6B and 6C). The MCC tree was compared with a CCD-0 tree and only minor changes in the topology were observed.

**Figure 6.**
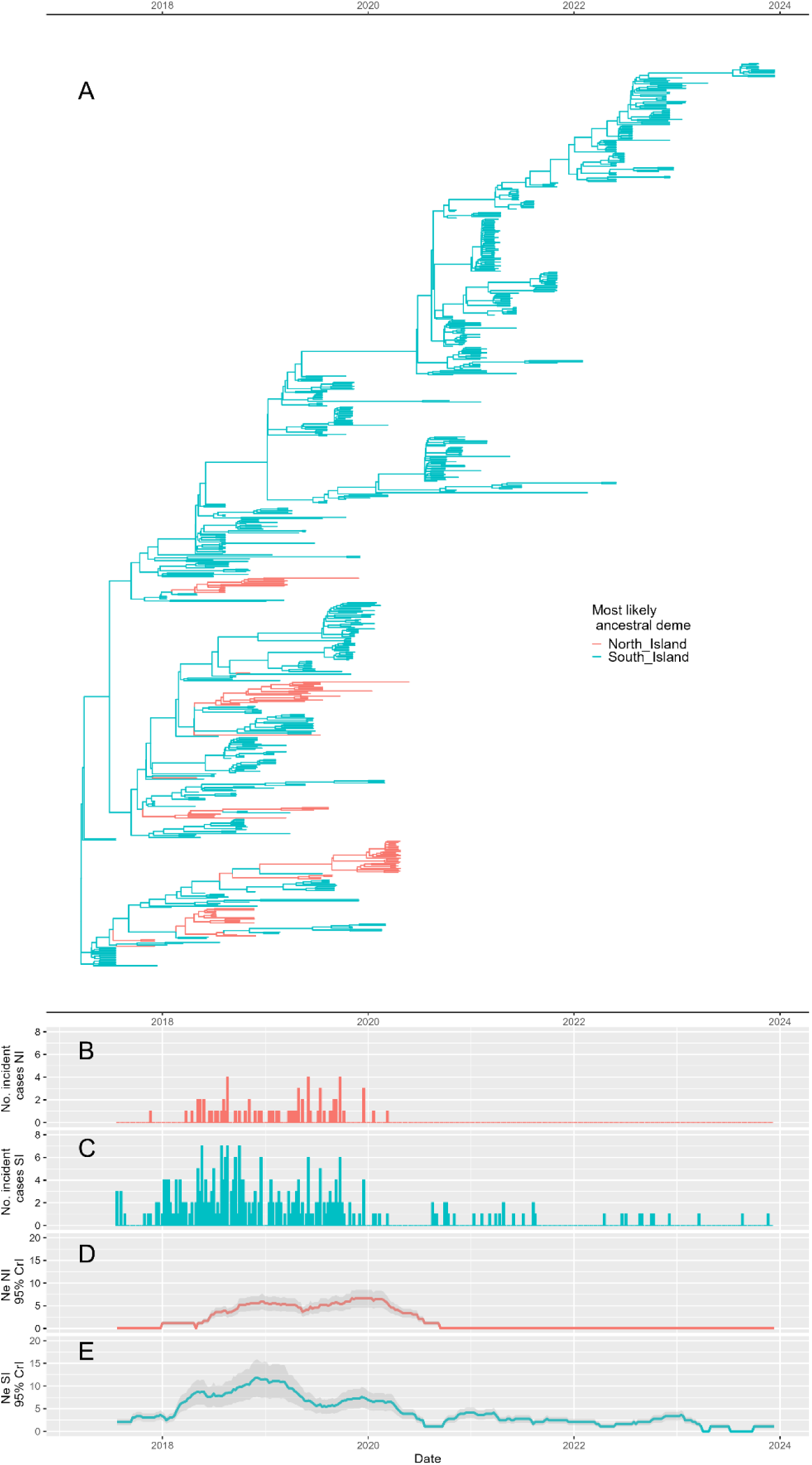
A: MCC tree from the two-deme MASCOT GLM analysis of the animal-level dataset of *M. bovis* isolates from New Zealand’s North and South Islands. Branches are coloured according to inferred ancestry. B and C: Number of confirmed infected farms (incidents per week) in the North Island (B) and the South Island (C). D and E: Effective population sizes, N_e_, inferred for the North Island (D) and the South Island (E), with shaded 95% HPD.

Transmission events involving the feedlot were examined using a 3-deme MASCOT-GLM whereby the feedlot was considered as a separate deme from the rest of the South Island (Supplementary Figures S4 and S5. The MASCOT-GLM analysis further supports the evidence from the SCOTTI model of multiple transmission events from the feedlot to other properties in the South Island.

## Discussion

Since it was first named in 2004, phylodynamic modelling has been widely used to explore the evolution, population dynamics and transmission of both viral and bacterial pathogens [51].

Phylodynamic studies have spanned large and small scales in both space and time, and include examples of globally circulating pathogens over relatively long [24, 52] and short [53] time scales and more localised pathogen dynamics [54, 55]. However, there are relatively few examples of the application of genomics and phylodynamics to inform decision making during an outbreak [56–59] and even fewer examples of the application of genomics and phylodynamics during outbreaks of animal pathogens [60–62].

This study presents a detailed phylodynamic analysis of an outbreak of the primary cattle pathogen *Mycoplasma bovis*, from its first reported incursion in 2017 to successful control and progress towards eradication in New Zealand. Although the work presented is largely a retrospective analysis of all data collected over the six years of the eradication programme, the tools described were used throughout the outbreak to provide unique insights into patterns of transmission and to support decision making by the programme. These were augmented by visualisation tools that allow interactive analyses of spatial, temporal, genomic and epidemiological data, including Nextstrain [20] and Microreact [21].

Evidence from a global analysis of *M. bovis* genomes by Binney, Gias (18) supported a single introduction of ST21 followed by evolution within New Zealand. The global analysis showed that New Zealand isolates formed a monophylectic clade, with evidence of multiple 7-gene MLST single locus variants evolving from ST21 within New Zealand [18]. Analyses of the tMRCA throughout the outbreak provided evidence for a single incursion that was relatively recent and unlikely to be more than two-years prior to the first detection in 2017. However, given the early divergence into three lineages the possibility of up to three incursions from the same international source cannot be ruled out. Several potential sources of infection have been considered including imported bovine semen, embryos, live cattle, other live animals, feed and equipment [63].

The analysis presented shows evidence of both long range and local transmission, supporting tracing and other epidemiological analyses. A detailed analysis of stock movement data revealed extensive movements of cattle across New Zealand between 2017 and 2020, with over 450,000 consignments transported per year [64]. The movements were highly seasonal, with the greatest in May and June each year, reflecting the concentration of movements of dairy cattle around the national ‘Moving Day’ on 31 May. The MASCOT GLM model indicated transmission between the two islands occurred on multiple occasions up to 2019. After that time no further ‘migrations’ between the islands were observed, and in 2021 active infection became confined to the Canterbury region in the South Island. Network analysis of the cattle movement data showed evidence of at least four major ‘communities’ that were largely geographically clustered, with three in the North Island and one in the South Island [64]. The South Island-associated ‘community’ extended into the North Island, particularly the southern region of the North Island, revealing connectedness between the two islands due to cattle movements. The community analysis showed that, while most movements were within islands, and within geographically defined communities, there was extensive movement of cattle consignments between communities located in different islands. For example in 2020 there were over 3,000 movements of cattle consignments from the South to the North Island and over 700 movements from the North to the South Island [64]. This is consistent with phylodynamic modelling estimates of multiple transmission events (median of 16) between the two islands, with more events from South to North than North to South.

The change in the effective reproduction number, R_eff_, over time estimated from the phylodynamic model was consistent with the estimates based on a branching process model of case data reported by Jordan, Sadler (8). Both studies report values greater than 2.0 prior to early 2018, when a sharp decline to less than 1.0 is observed. This coincides with the commencement of the eradication programme on 28 May 2018, following the initial detection in July 2017. Jordan, Sadler (8) estimated R_eff_ using data available up to November 2019, when the peak of transmission was over and there were few active infected farms. The present study extended these estimates to the end of 2023 and provided evidence of a small increase in R_eff_ after 2021 prior to the absence of known infection at the time of writing.

The effective population size estimates, N_e_, in the context of this analysis are an estimate of the population of infected individuals at one time point that give rise to the next generation of infected individuals and can indicate the dynamics of infection between animals and farms. For example, an increase in N_e_ may indicate a population of infected individuals that is growing because of transmission, and a decline may indicate an evolutionary bottleneck due to the elimination of infections. The BICEPS and MASCOT GLM models show a similar trend in N_e_, peaking in 2018/19, declining after the onset of the eradication programme but with evidence of increases in transmission during 2021 when infection was localised in the Canterbury region.

The ‘long tail’ of localised infection in the Canterbury region was a striking feature of the New Zealand *M. bovis* outbreak. A similar phenomenon was observed in the 2001 Foot and Mouth Disease virus (FMDv) outbreak in the United Kingdom. The tail in the UK FMDv outbreak was localised in the Cumbria region of the UK and attributed to within-farm transmission [65]. Similarly, within-farm transmission with local spillover of infection may have caused the persistence of infection with *M. bovis* in New Zealand. A large feedlot in the Canterbury region was detected as infected in August 2018, and this farm was not depopulated until December 2022. It was initially considered unlikely that this farm was a source of infection to other farms due to the farm operating as a receiver of cattle destined for slaughter. However, the observation of multiple farms in the proximity of the feedlot becoming infected, and the genomic evidence of multiple chains of transmission with distinct lineages common to the feedlot, challenged the assumption that the feedlot was solely a ‘sink’ for transmission, and instead may have been a source of infection to other herds [16, 66]. For the first time in the programme, a Controlled Area Notice was placed in the Canterbury region surrounding the feedlot in October 2022 prior to depopulation of the feedlot. These additional measures facilitated the elimination of infection in the area [66].

Phylodynamic modelling using the SCOTTI model provided additional evidence that the feedlot was a source of infection for local farms. Further evidence for outward transmission from the feedlot was provided by the MASCOT GLM model, which estimated over 40 transmission events from the feedlot to other South Island farms. The number of events and direction of transmission may be biased by the relatively high proportion of isolates in the animal-level dataset that were from infected cattle on the feedlot sampled over the four-year period (14%, 99/697). Further, the estimated size and direction of transmission estimated by the SCOTTI model is informed by both the genomic signal and the period of infectiousness for farms in the transmission network. The lower bound of the period of infectiousness is informed by the estimated date at which farms became infected, which was often uncertain. Although varying the infection dates did not affect the overall conclusion, uncertainty in the infectious period and the higher proportion of isolates from the feedlot may have biased estimates of both the direction and size of transmission probabilities. Despite the potential bias in the direction of transmission estimated by the SCOTTI model, further investigation into other routes of transmission was warranted, including the possibility of unrecorded cattle movements [64], airborne spread [4], and transmission via fomites [11].

Epidemiological data and phylodynamic modelling supported the implementation of the New Zealand *M. bovis* eradication programme in 2018. The extensive application of genome sequencing provided an additional layer of information to determine the most likely time of initial incursion, and patterns of transmission. New cases were often identified through tracing cattle movements, indicating this was the most important transmission pathway between farms. However, new cases may be linked to multiple case farms through movement tracing, making it impossible to determine the source of infection using movement data alone [8]. In these situations, genomic data and phylodynamic analysis helped to identify the most likely transmission events identified through movement tracing. However, there were examples where inferences from phylodynamic models did not agree with those from farm-level epidemiological data, including tracing data, particularly when there were multiple potential infection sources. This highlighted the importance of taking an integrative approach whereby genomic analyses, using a range of phylodynamic models, complemented conventional epidemiological approaches. This was most notable during the outbreak when different models provided consistent support for hypotheses concerning the source of new infections, containment of the outbreak and the absence of new incursions. Assigning infected farms to clades allowed prioritisation of investigative work to extant lineages. For example, a review of formerly infected farms assessed if new information revealed risk that was not covered at the time of initial investigation. Often farmers would be contacted to validate information such as cattle movements. Instead of reviewing all 282 farms, work was focussed on those associated with recently circulating lineages. This was an efficient way to gather confidence of absence and minimise disruption to farmers who may have otherwise been contacted by the programme.

New Zealand is the first country to attempt to eradicate *M. bovis* and there is optimism that the Government and industry-led programme has been successful. Genomics and phylodynamic modelling have contributed to the programme by complementing other epidemiological analyses as the outbreak progressed. In 2022, genomic analysis also confirmed that a dairy farm in the Canterbury region was infected with a different ST (ST29) following a bulk milk detection [16, 66]. This was confined to the single farm, and it was determined that the most likely source was semen imported into the country prior to implementation of the current Import Health Standards for bovine germplasm in May 2022 [66]. Therefore, to date there is a strong indication that both the eradication programme and the measures taken to prevent re-introduction have been effective, underlining the feasibility of such a programme in an island nation with a large cattle industry.

## Acknowledgements

Simon Firestone’s contribution to this research was supported by the Australian Government through the Australian Research Council’s Discovery Projects funding scheme (project DP210103239).

## Author contributions

All authors contributed to writing the manuscript (original draft preparation, review and editing). NF, DW and PB contributed to conceptualisation, formal analysis, data curation and methodology. SF, SB and BB contributed to formal analysis and methodology. JF contributed to methodology and resources. AB and MvA contributed to resources, project administration and funding.

---

**Table S1.**
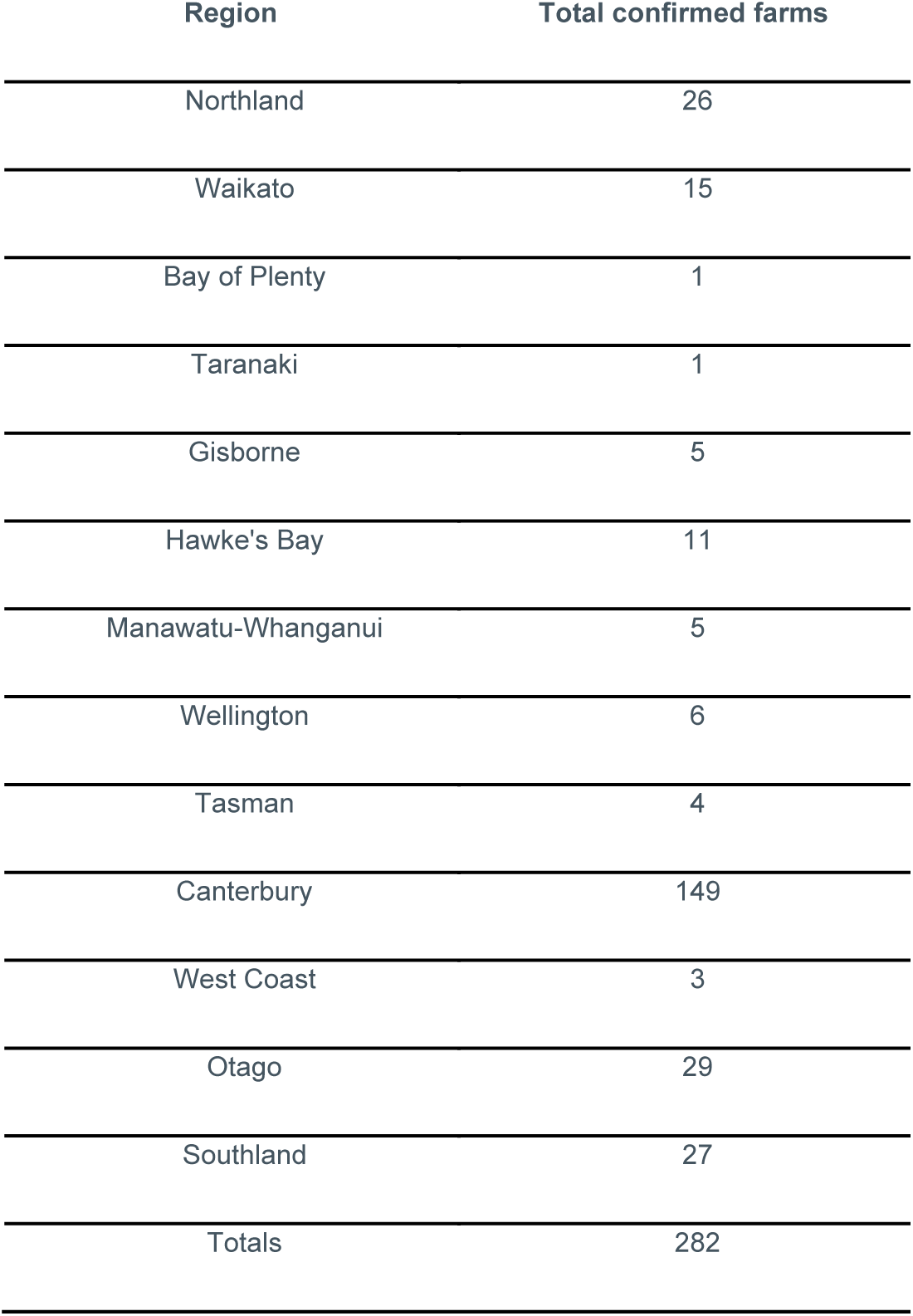
The number of farms confirmed as *Mycoplasma bovis* positive by region of New Zealand.

**Supplementary Figure S1.**
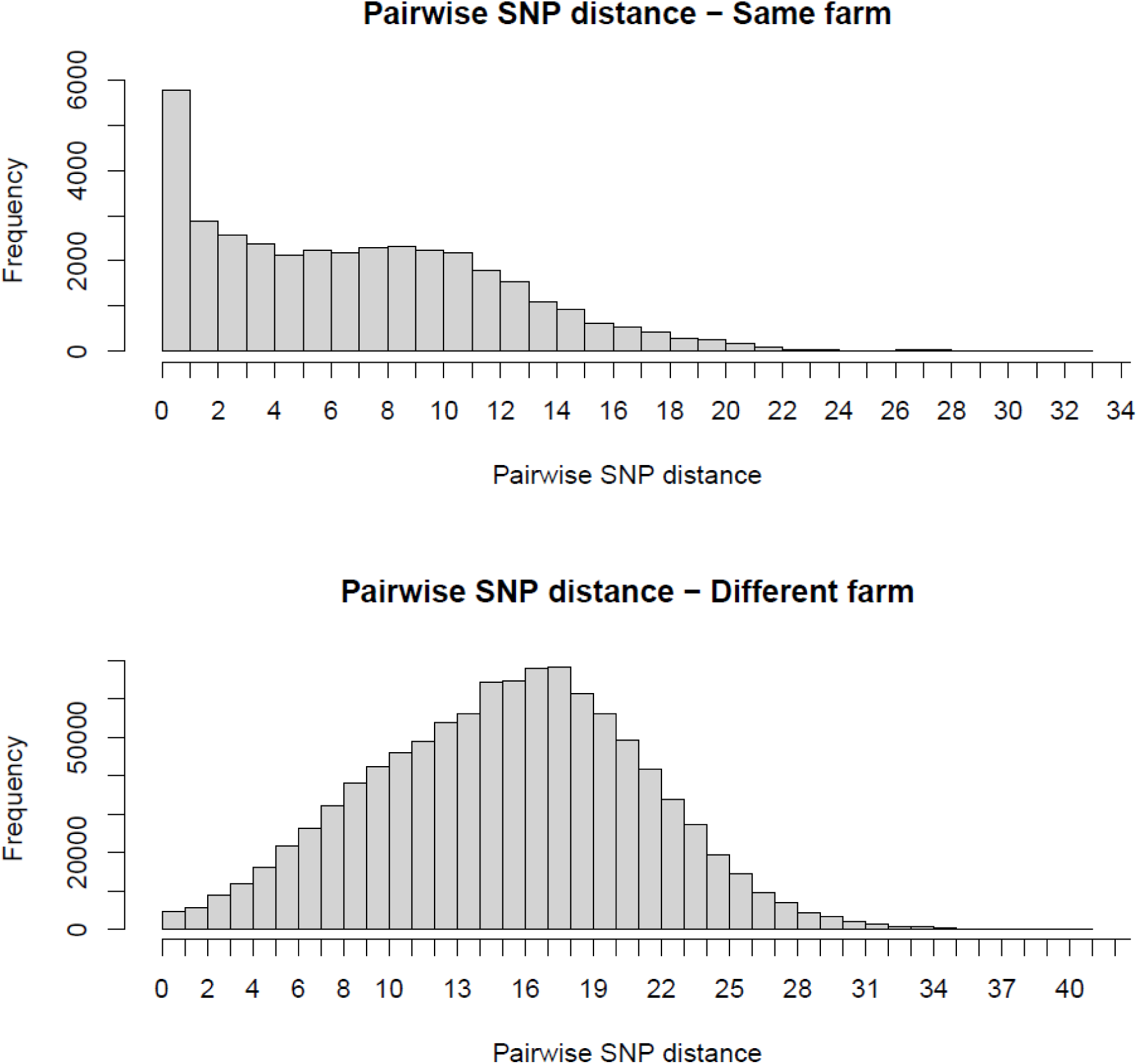
Pairwise single nucleotide polymorphism differences between 697 *Mycoplasma bovis* isolates from the same farm and different farms in the New Zealand outbreak.

**Supplementary Figure S2.**
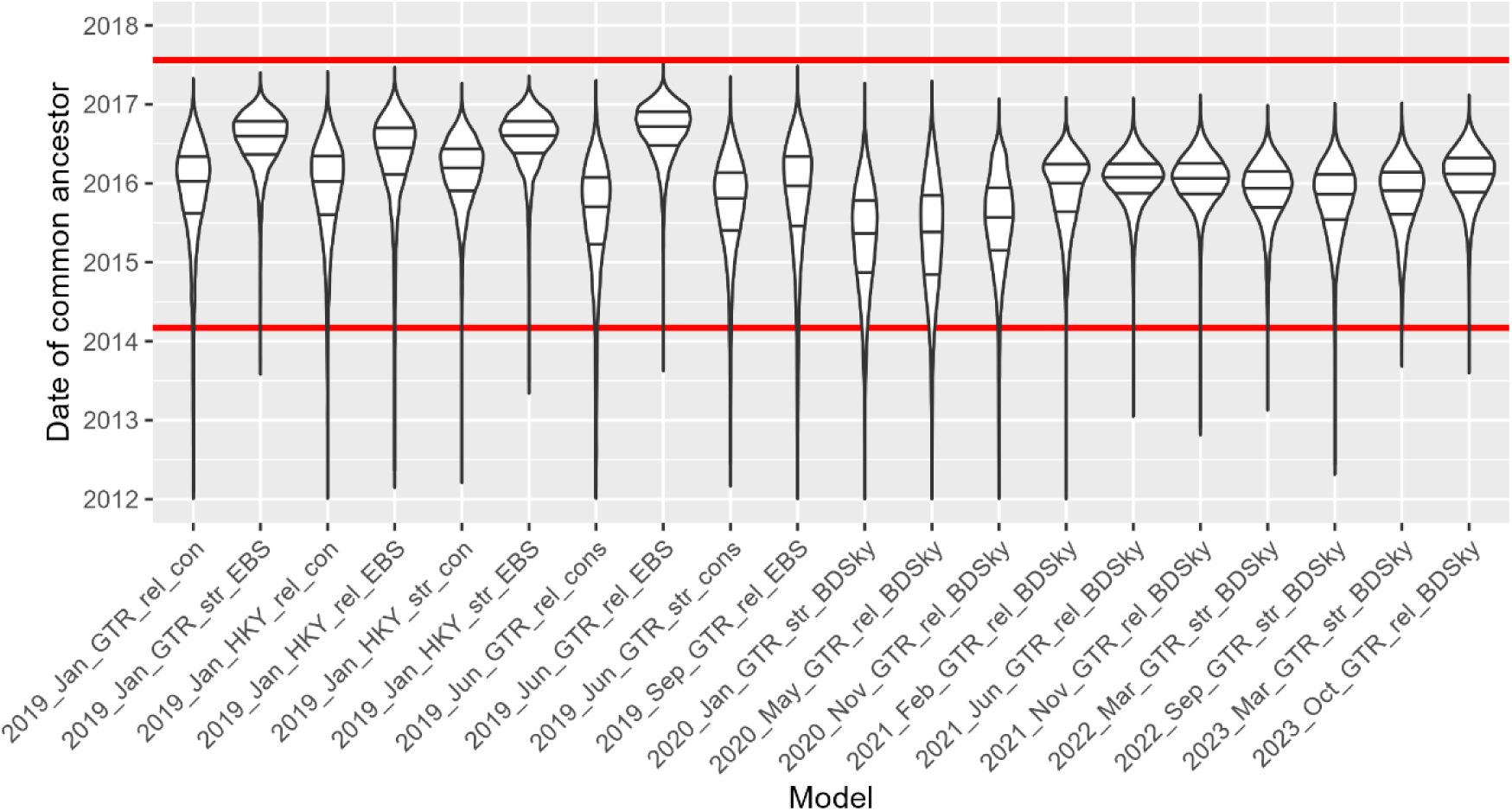
Estimated dates of common ancestor of isolates collected from *Mycoplasma bovis* infected farms between January 2019 and October 2023. The different models are captured in the column labels. Substitution models were Generalised Time Reversible (GTR) or Hasegawa-Kishino-Yano (HKY) models, clock models were either strict (str) or relative (rel) and tree models were either constant coalescent (cons), Extended Bayesian Skyline (EBS) or Birth-Death Serial Skyline (BDSky) models. The horizontal lines within the violin plots show the median and interquartile ranges. The upper red line shows the date the first case was detected, and the lower red line shows the earliest hypothesised infection date from epidemiological data. The choice of model was informed by nested sampling.

**Supplementary Figure S3.**
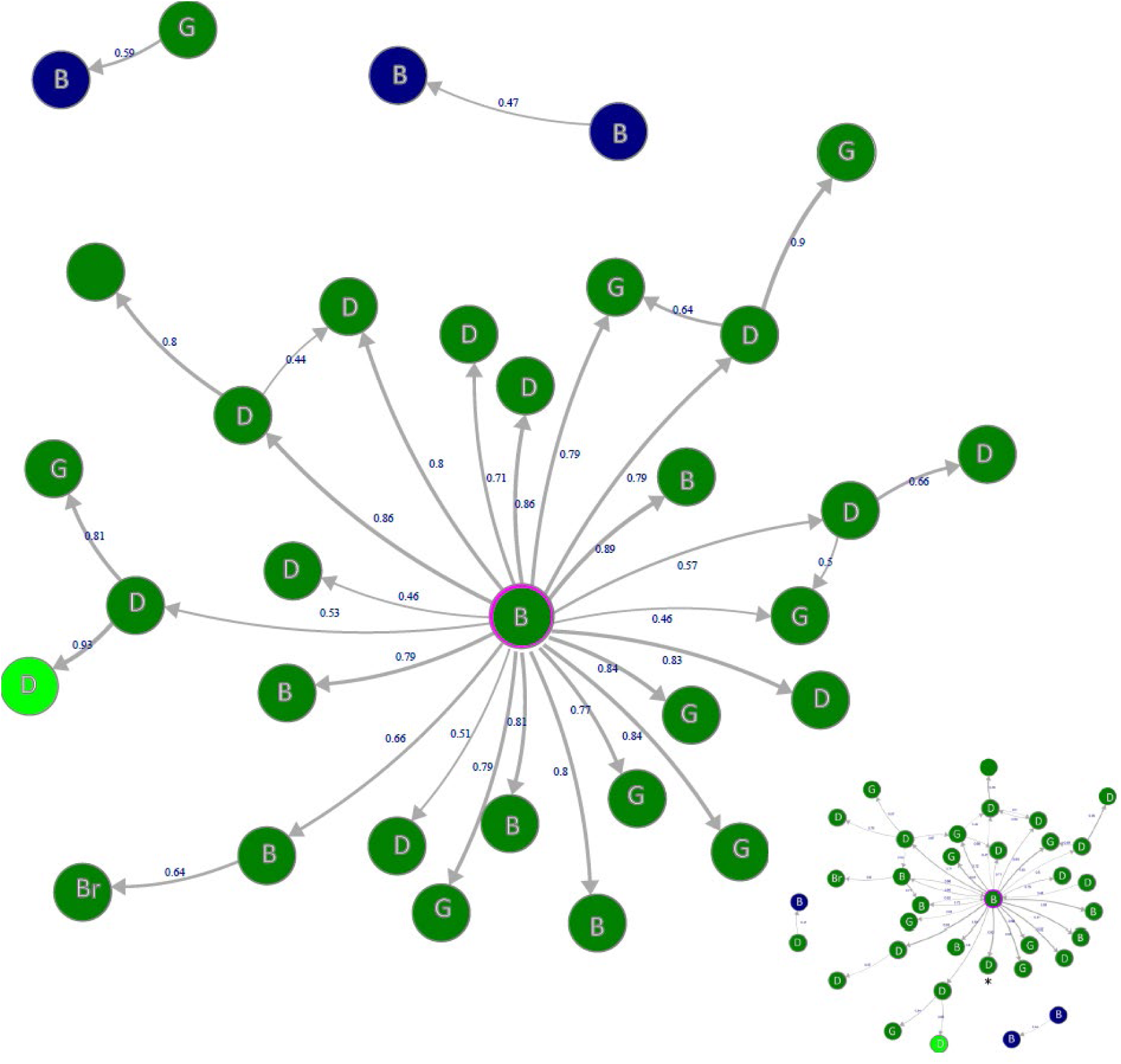
Network diagrams showing between-farm direct transmission probabilities estimated using the SCOTTI model, using all lineage ‘a’ isolates (RB levels 2, 4 and 5) up to December 2023 and a date of onset of infectiousness of one month prior to the imposition of movement restrictions (main figure). The inset is Figure 4B for comparison.

**Supplementary Figure S4.**
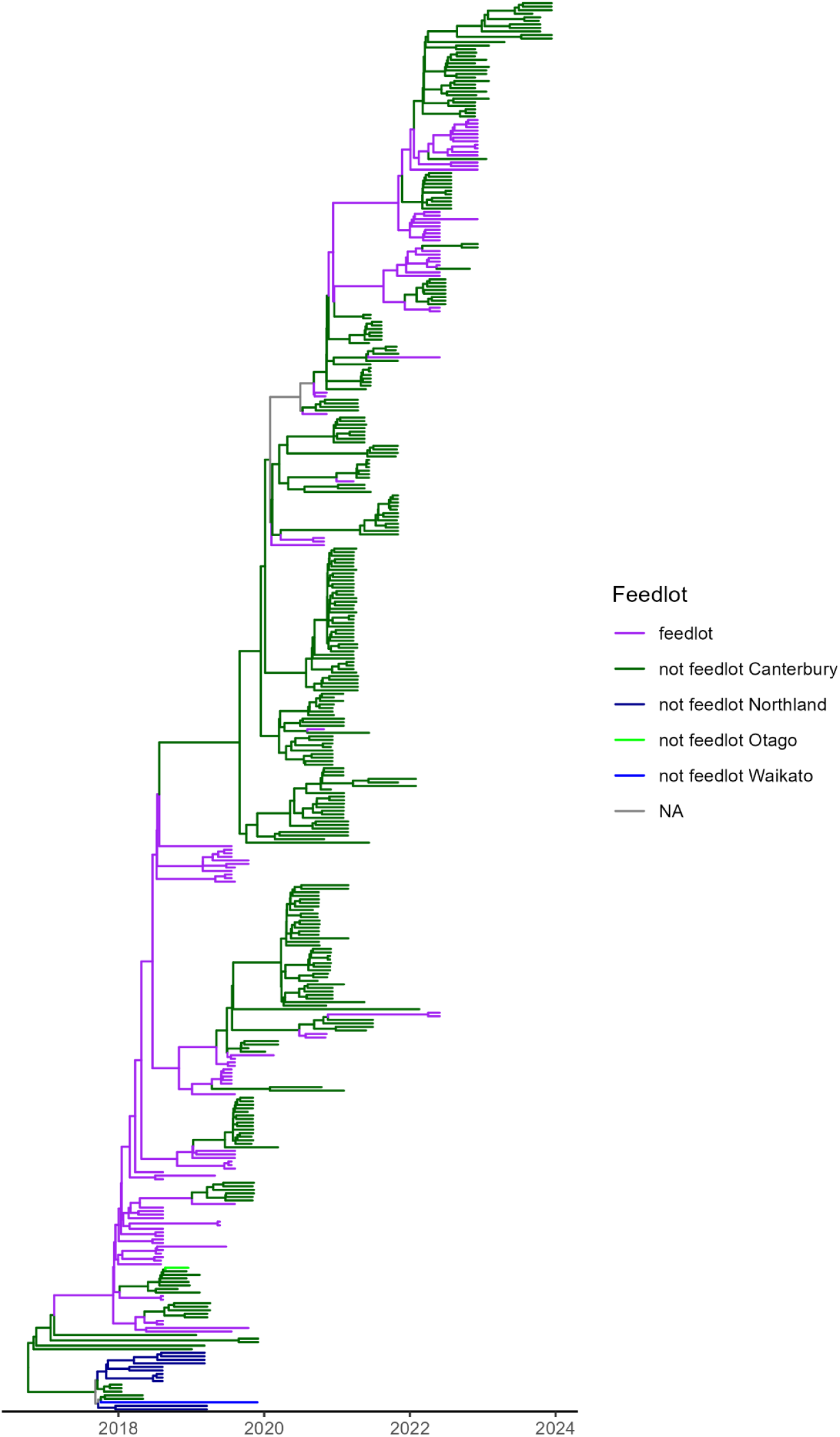
Consensus tree from the SCOTTI analysis of *Mycoplasma bovis* lineage ‘a’ in New Zealand. The large feedlot is shown in purple, indicating evidence of ancestral states assigned to this farm.

**Supplementary Figure S5.**
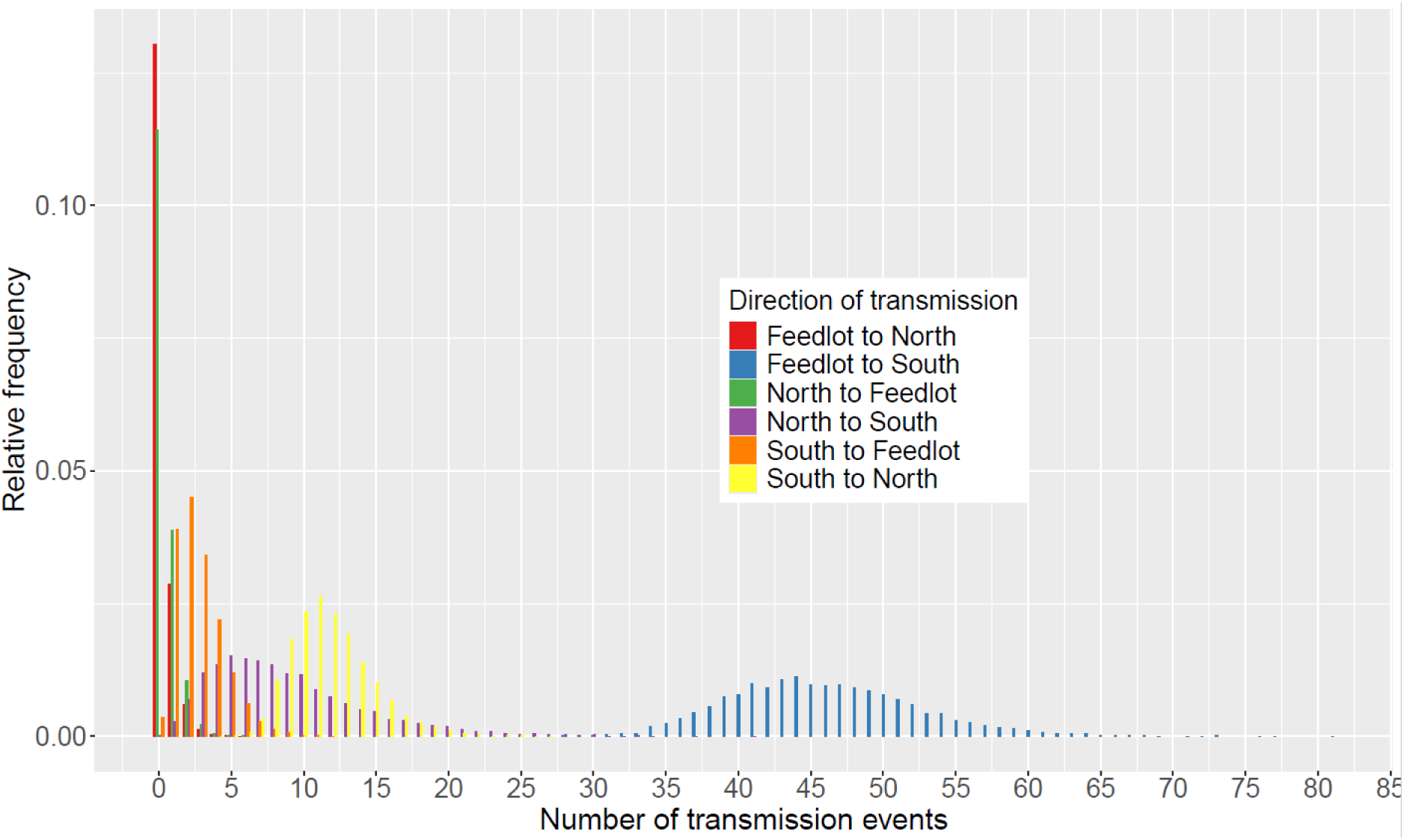
Posterior distributions of estimated between-island *M. bovis* transmission events from a 3 deme marginal structured coalescent approximation model, MASCOT GLM [48, 49], in BEAST2 [22]. In this analysis the feedlot was considered as a separate deme from the rest of the South Island.

